# Sexual selection in females across the animal tree of life

**DOI:** 10.1101/2021.05.25.445581

**Authors:** Salomé Fromonteil, Lennart Winkler, Lucas Marie-Orleach, Tim Janicke

## Abstract

The pioneers of sexual selection theory proposed that males are generally ‘eager’ whereas females are rather ‘coy’ with respect to mating. This male-centred perspective on sexual selection continues to permeate our perception of sex differences across disciplines. Despite an increased awareness that females also compete for mating partners, we still tend to consider sexual selection in females a rare peculiarity. Here we present meta-analytic evidence from 72 species across a broad range of animal taxa to show that sexual selection in females is widespread and should be considered the norm rather than the exception. Thereby, our results extend our general understanding of sexual reproduction and may contribute to a more balanced perspective of how sexual selection operates in both males and females.

## Introduction

Darwin’s original conception of sexual selection came along with the assertion that it is primarily the male that competes for access to mates whereas the female chooses among mates. Darwin argued that this sex difference is virtually ubiquitous, claiming that “with almost all animals, in which the sexes are separate, there is a constantly recurrent struggle between the males for the possession of the females” and that “the female […], with the rarest exception, is less eager than the male […,] she is coy and may often be seen endeavouring for a long time to escape from the male” (*1*). Ever since, this stereotypic perspective of eager males and reluctant females continues to predominate the field of sexual selection and presumably contributed to a pervasive bias in research agendas of behavioural ecologists and evolutionary biologists over the last decades (*2*–*4*). Until now, studies exploring aspects of sexual selection on males in terms of male-male competition and female choice massively outnumber those examining female competition for mates and male choice by a magnitude of ten (Figure S1).

Notwithstanding the long-lasting bias in our efforts to understand male and female reproductive strategies, there is both theoretical and empirical support for stronger sexual selection on males. In particular, Bateman’s milestone contribution paved the way towards a better understanding of the evolutionary causes of sex differences (*5*). He argued that reproductive success of males but not females is primarily governed by the number of mating partners, which is ultimately rooted in anisogamy and gives rise to prevalent intra-male competition. The postulated higher benefit of an additional mating in males has been found to prevail in the animal kingdom (*6*) and various lines of theoretical work support the view that the primordial sex difference in gamete size promotes stronger sexual selection on males (*7*–*9*).

The predominance of stronger competition among males and choosiness in females is often accompanied by female-biased parental care (i.e., post-zygotic investment; (*10*, *11*)) — a behavioural syndrome that has been termed “conventional” or “Darwinian” sex role (*6*, *12*). However, the usefulness of this sex-role concept is controversial until today, especially because it may prevent us from exploring and acknowledging the entire spectrum of sex differences in nature in an unbiased manner (*13*–*16*). Eventually the question arises whether evidence for sexual selection being typically stronger in males necessarily implies that sexual selection is rare in females. In fact, there are a number of illustrative examples for female-female competition and male choice at both pre- and post-copulatory episodes of sexual selection suggesting that it can act on females in a very similar way as it does on males (*17*–*20*). The most prominent and clearest support for sexual selection in females can be found in so-called sex-role reversed species in which females often compete actively for males and are the more ornamented sex. For instance, in some species of pipefishes and seahorses, fertilization takes place inside the brood pouch of the male, which provides all parental care (*21*, *22*). As a consequence, males become a limiting resource for which females compete, which eventually leads to selection for ornaments favoured by male pre- and even post-copulatory mate choice (*23*). Other illustrative examples for sex-role reversal are tropical shorebirds of the family *Jacanidae* in which females aggressively defend territories to monopolize multiple males (*24*). Importantly however, sex-role reversal is not a prerequisite for sexual selection to operate in females, as it may represent just one extreme on a spectrum of sex-roles. Even in species with conventional sex roles in which sexual selection promotes the evolution of male ornaments and extravagant courtship behaviours, females may still compete for access to high-quality males, as demonstrated in male lekking fruit flies (*25*) and peafowls (*26*). Consequently, sexual selection in females might actually be an omnipresent phenomenon in animals but operating less intensely and more subtly compared to males, which can make it more difficult to detect.

Considering the increased awareness that competition for mates also occurs in females, the critical question remains whether all qualitative examples represent exceptions from the general rule or whether sexual selection is actually a widespread evolutionary force in females too. Here we present a meta-analytic approach to inform the ongoing debate on sex roles by providing a quantitative test for female sexual selection across the animal tree of life. We compiled published estimates of the so-called Bateman gradient, which measures the fitness benefit of mating. Hence, this metric captures the selective advantage arising from intra-sexual competition for mates, which is the essence of Darwinian sexual selection (*1*, *5*) and therefore constitutes a powerful measure to compare the strength of sexual selection across a broad array of species (*27*, *28*). We ran a series of phylogenetically controlled analyses showing that (*i*) sexual selection on females is the norm rather than the exception, (*ii*) positive female Bateman gradients are robust to varying methodological approaches of primary studies, and (*iii*) a higher benefit of mating in females translates into a more polyandrous mating system.

## Methods

### Systematic literature search

We extracted female Bateman gradients from a previous meta-analysis (*6*) and expanded this database by adding studies that have since been published. Specifically, we ran a systematic literature search using the ISI Web of Knowledge (ISI Web of Science Core Collection database; Clarivate Analytics) with the ‘topic’ search terms defined as (“Bateman*” OR “opportunit* for selection” OR “opportunit* for sexual selection” OR “selection gradient*” OR (“mating success” AND “female*”)) on the 30^th^ of November 2020. In this search, the timespan was defined as “2015 – today” because the literature search of the previous study had been carried out on the 25th of April 2015. In addition, we also screened all studies published after 2015 that cited Bateman’s original paper. Our sole inclusion criterion was that the study must report data allowing to assess the relationship between mating success and reproductive success for females. The additional search yielded 2,140 records of which 26 studies were considered eligible, providing a total of 35 new estimates of female Bateman gradients. In addition, we included 4 estimates from an unpublished experimental study on the bean weevil *Acanthoscelides obtectus* (S. Fromonteil, unpublished data). Combining these estimates with the ones obtained from the previous meta-analysis added up to a final dataset of 78 studies reporting 111 female Bateman gradients from 72 species (Figure S2).

### Moderator variables

Apart from a global test of sexual selection in females (inferred from a positive Bateman gradient), we aimed at explaining among-study variation in effect sizes from both a methodological and an evolutionary perspective. First, we evaluated if the method to quantify mating success influenced Bateman gradients. Especially for females, the measurement of mating success in terms of the number of genetic parents (i.e., genetic mating success) has been demonstrated repeatedly to overestimate the Bateman gradient when compared to estimates obtained from behavioural observations (i.e., copulatory mating success) (*27*). Quantifying mating success in terms of the number of genetic parents may not only obscure a potentially important component of post-copulatory sexual selection (because unsuccessful copulations and multiple copulations with the same partner remain undetected) but also leads to an autocorrelation of mating success and reproductive success, particularly in species with low fecundity (*29*). For those reasons, we tested the effect of the mating success method by contrasting estimates of Bateman gradients based on genetic (*k* = 74) *versus* copulatory mating success (*k* = 37). Second, we explored the impact of having unmated individuals included in the measurement of the Bateman gradient. Estimates including this zero-mating success category provide a combined estimate for the benefit of mating once and the benefit of having an additional mating partner (or copulation), whereas Bateman gradients excluding zero-mating success data capture only the latter. In the context of sexual selection, we are primarily interested in the benefit of having an additional mating partner (or copulation) rather than the benefit of mating itself, since the latter is essential for reproduction in outcrossing species and therefore rather trivial. Thus, we compared Bateman gradients that include unmated individuals (*k* = 61) with those excluding this zero-mating success category (*k* = 71). Third, to further account for methodological differences between studies, we tested for an effect of the study type on Bateman gradients by comparing field studies (*k* = 65) with laboratory studies (*k* = 46).

Fourth, we tested whether the Bateman gradient was related to the mating system. We predicted that a fitness benefit of achieving a higher mating success selects for increased polyandry, meaning that species with steeper Bateman gradients are expected to be more polyandrous (*30*). We classified the mating system of each sampled species based on estimates of polyandry, which we defined as the proportion of reproducing females that have more than one mating partner. For the majority of species (*N* = 61; 84.7%), we estimated the proportion of multiply mated females using data provided in the primary studies (Table S3). For most of the remaining species, we extracted estimates of polyandry from secondary literature, except for three species for which we could only find verbal classifications of the mating system (see Table S3 for references). We then used these estimates to define the mating system as either monandrous or polyandrous, depending on whether its value was lower or higher than 0.5, respectively, because this value has been found to be the average level of polyandry in wild populations (*31*). Hence, species classified as monandrous are not strictly monandrous in the sense of having only one mating partner. Instead, monandrous species also mate multiply but at a lower frequency compared to polyandrous species. In total, our dataset encompassed 16 monandrous and 56 polyandrous species, for which we obtained 32 and 79 effect sizes, respectively. Sensitivity analyses revealed that alternative thresholds of polyandry (i.e., 0.4 or 0.3) did not lead to qualitative changes of results. We note that our classification of the mating system remains an oversimplification of a clearly more gradual spectrum of natural mating systems. For this reason, we also ran an alternative model in which we used the actual estimate of polyandry as a continuous predictor variable instead of the explanatory factor mating system.

### Phylogenetic affinities

We reconstructed the phylogeny of all sampled species from published data in order to account for phylogenetic non-independence (Figure S3). Specifically, we extracted divergence times from the TimeTree database (http://www.timetree.org/; (*32*) and transformed the distance matrix into the NEWICK format using the unweighted pair group method with arithmetic mean (UPGMA) algorithm implemented in MEGA (https://www.megasoftware.net/; (*33*). In total, our analysis included 72 species with a broad distribution across the animal tree of life, with overrepresentation of arthropods (*N*_Species_ = 19), birds (*N*_Species_ = 14), fishes (*N*_Species_ = 13), and mammals (*N*_Species_ = 8) (Figure S3).

### Statistical analysis

The Bateman gradient is defined as the slope of a linear regression of reproductive success on mating success (*5*) and provides a powerful metric of the strength of sexual selection for inter-specific comparisons when computed on relativised data (i.e., accounting for differences in mean mating and reproductive success) (*34*). However, only 57.6 % of the extracted Bateman gradients were computed on relativised data. Therefore, we converted all obtained slopes into Pearson correlation coefficients (*r*) and computed their sampling variances using formulas reported elsewhere (*35*) (Figure S4). We note that using *r* as an effect size instead of a slope quantifies the strength of the relationship between mating success and reproductive success, which depends not only on the slope (i.e., the fitness return of the mating) but also on the goodness of fit (i.e., the standard error of the slope). However, analysis of the subset of data for which we have standardised Bateman gradients revealed that *r* is a strong predictor of the actual Bateman gradient (Linear Regression: estimate ± SE = 1.16 ± 0.06; *F*_1,66_ = 322.21; *P* < 0.001, *R*^2^ = 0.84), suggesting that our effect size is a reliable estimate for the benefit of mating.

We ran General Linear Mixed-Effects Models (GLMMs) to provide a global test for sexual selection in females and to explore determinants of the inter-study variation. First, we quantified global effect sizes by running GLMMs with *r* defined as the response variable weighted by the inverse of its sampling variance, and included study identifier and observation identifier as a random term. This was done both without (i.e., ‘non-phylogenetic’ GLMMs) and with adding the phylogenetic correlation matrix as an additional random term (‘phylogenetic’ GLMMs). Secondly, we ran phylogenetic GLMMs in which we defined mating success method (copulatory *versus* genetic), mating success range (with *versus* without zero-mating success category), study type (field *versus* laboratory studies), or mating system as a fixed factor to explain inter-study variation in *r*. In order to complement our analysis of the mating system, we also ran a phylogenetic GLMM including estimates of the actual level of polyandry (i.e., the proportion of multiply mated females) as a continuous predictor variable. All GLMMs were run with the MCMCglmm function of the MCMCglmm R package version 2.29 (*36*), using uninformative priors (*V* = 1, *nu* = 0.002) and an effective sample size of 10,000 (number of iterations = 4,400,000, burn-in = 400,000, thinning interval = 400). All models were also run with alternative priors, which revealed qualitatively identical results. Moreover, we ran all models multiple times to verify convergence and checked for autocorrelation in the chains. For completeness, we also ran all GLMMs using the Restricted Maximum Likelihood (REML) approach using the metafor R package version 2.4-0 (*37*). These complementary analyses provided qualitatively similar results and are reported in the Supplementary Material (Table S1 and S2).

We estimated heterogeneity *I*^2^ from the intercept-only model as the proportion of variance in effect size that can be attributed to the different levels of random effects (*38*). In particular, we decomposed total heterogeneity into the proportional phylogenetic variance (*I*^2^_Phylogeny_), between-study variance (*I*^2^_Study_), and study-specific variance (observation-level random effect; *I*^2^_Observation_) (*39*). Note that *I*^2^_Phylogeny_ is also termed phylogenetic heritability *H*^2^ and is equivalent to Pagel’s *λ* (*40*). For models including predictor variables, we computed the proportion of variance explained by those fixed factors (‘marginal *R*^2^’) (*41*).

We used Kendell’s rank correlation test to quantify funnel plot asymmetry, which can be indicative of publication bias. Moreover, we tested whether the year of publication influences effect sizes, which has been argued to be suggestive of other forms of biases (*42*). For instance, the so-called bandwagon effect suggests that supportive results get easier published in a newly emerging field but over time scepticism about the theoretical foundations may arise and initially non-intuitive findings may find a more receptive audience. If true for the field of sexual selection, we may expect an increase of effect sizes for female Bateman gradients with the rising awareness in the community that sexual selection does not only operate in males.

All statistical analyses were carried out in R version 4.0.3 (*43*).

## Results

We found evidence for sexual selection in females to be common across the animal tree of life inferred from a positive global effect size of the Bateman gradient (Figure 1A; Table 1 & S1). However, estimates of sexual selection showed substantial heterogeneity across studies (Figure S4; Table 1). This variation could partly be explained by differences in methodological approaches used to quantify the strength of sexual selection. Specifically, estimates of sexual selection critically depended on how mating success was measured (Table 2 and S2; Figure S5): higher effect sizes were observed in studies using genetic parentage analysis to assess mating success (i.e., genetic mating success) compared to estimates based on behavioural observations (i.e., copulatory mating success). In addition, inclusion of individuals that did not mate, led to larger effect sizes compared to estimates excluding individuals with zero mating success (Table 2 and S2; Figure S5). However, we still observed a signal for positive selection on mating success when running more conservative analyses restricted to studies relying on copulatory mating success or excluding individuals that did not mate (Table 1 & S1). By contrast, we did not detect a significant difference in female Bateman gradients between laboratory and field studies (Table 1 & S1).

**Fig. 1.**
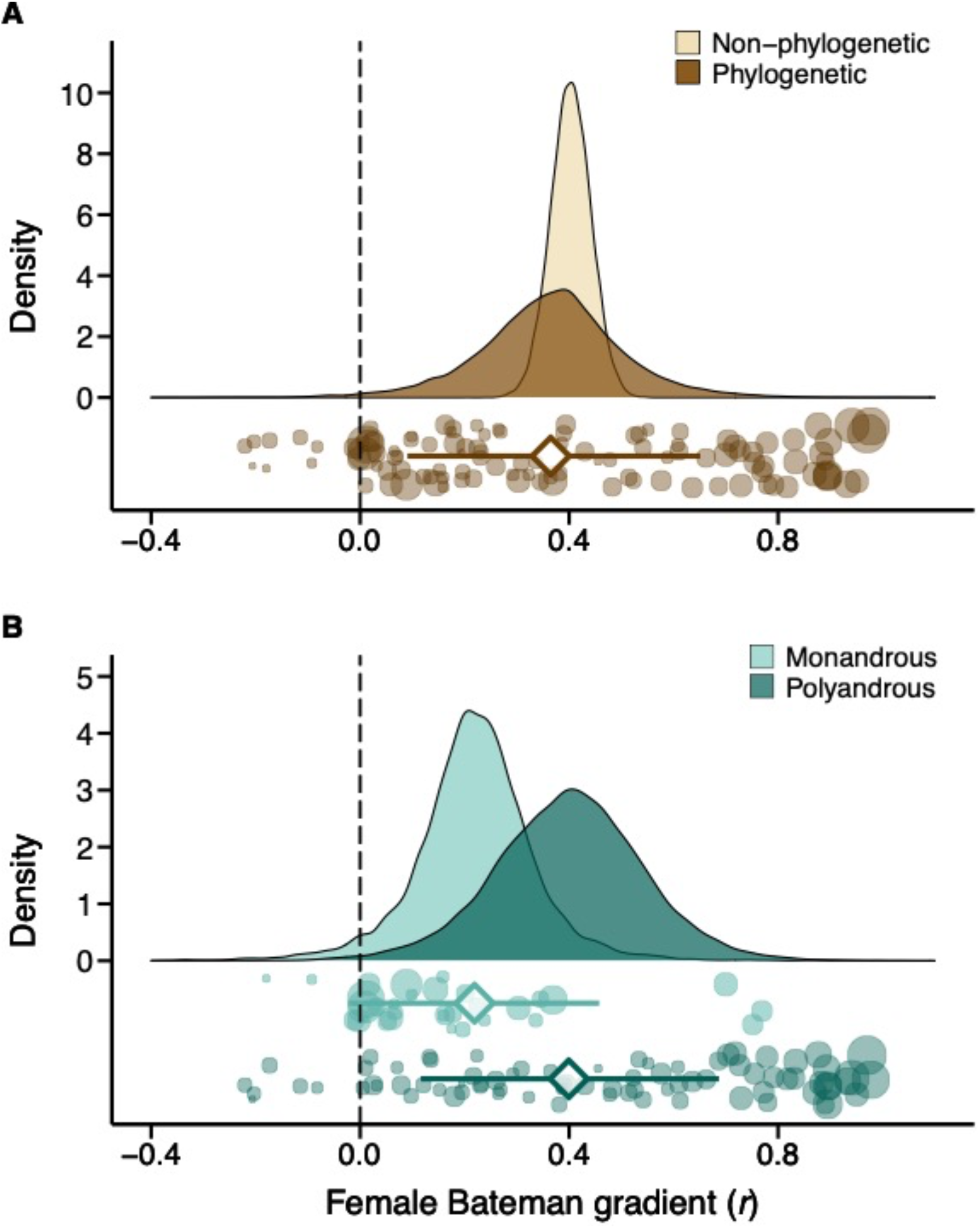
Meta-analytic evidence for sexual selection in females and its relation to the mating system. (A) Global effect size of the Bateman gradient obtained from Generalized Linear Mixed Models (GLMMs) with or without accounting for phylogenetic non-independence (phylogenetic or non-phylogenetic, respectively). (B) Contrast in sexual selection in females between monandrous and polyandrous species. Raincloud charts show posterior distributions, global effect size with 95% Highest Posterior Density intervals (diamonds and error bars) and raw effect sizes (filled circles) of female Bateman gradients.

**Table 1.**
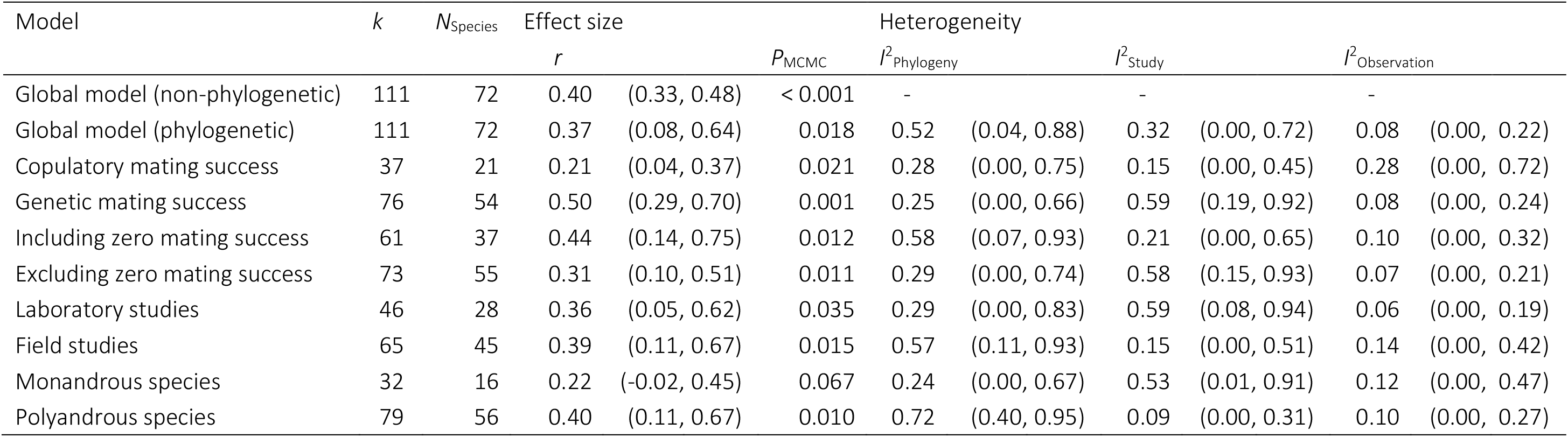
Global tests of sexual selection in females. Results of intercept-only phylogenetically controlled General Linear-Mixed Effects Models are shown for the entire dataset (global model) and subsets with respect to mating success method (copulatory *versus* genetic), mating success range (including *versus* excluding zero mating success category), study type (laboratory *versus* field studies) and mating system (monandrous *versus* polygamous species). Table shows number of effect sizes (*k*), number of species (*N*), effect size (*r*), and heterogeneity *I*^2^ arising from phylogenetic affinities, between-study variation, and between-observation variation. Model estimates are shown as posterior modes with 95% Highest Posterior Density (HPD) intervals.

| Model | *k* | *N*<sub>Species</sub> | Effect size |  | *P*<sub>MCMC</sub> | Heterogeneity |  |  |  |  |  |
| --- | --- | --- | --- | --- | --- | --- | --- | --- | --- | --- | --- |
|  |  |  | *r* |  |  | *I*<sup>2</sup><sub>Phylogeny</sub> |  | *I*<sup>2</sup><sub>Study</sub> |  | *I*<sup>2</sup><sub>Observation</sub> |  |
| Global model (non-phylogenetic) | 111 | 72 | 0.40 | (0.33, 0.48) | < 0.001 | - |  | - |  | - |  |
| Global model (phylogenetic) | 111 | 72 | 0.37 | (0.08, 0.64) | 0.018 | 0.52 | (0.04, 0.88) | 0.32 | (0.00, 0.72) | 0.08 | (0.00, 0.22) |
| Copulatory mating success | 37 | 21 | 0.21 | (0.04, 0.37) | 0.021 | 0.28 | (0.00, 0.75) | 0.15 | (0.00, 0.45) | 0.28 | (0.00, 0.72) |
| Genetic mating success | 76 | 54 | 0.50 | (0.29, 0.70) | 0.001 | 0.25 | (0.00, 0.66) | 0.59 | (0.19, 0.92) | 0.08 | (0.00, 0.24) |
| Including zero mating success | 61 | 37 | 0.44 | (0.14, 0.75) | 0.012 | 0.58 | (0.07, 0.93) | 0.21 | (0.00, 0.65) | 0.10 | (0.00, 0.32) |
| Excluding zero mating success | 73 | 55 | 0.31 | (0.10, 0.51) | 0.011 | 0.29 | (0.00, 0.74) | 0.58 | (0.15, 0.93) | 0.07 | (0.00, 0.21) |
| Laboratory studies | 46 | 28 | 0.36 | (0.05, 0.62) | 0.035 | 0.29 | (0.00, 0.83) | 0.59 | (0.08, 0.94) | 0.06 | (0.00, 0.19) |
| Field studies | 65 | 45 | 0.39 | (0.11, 0.67) | 0.015 | 0.57 | (0.11, 0.93) | 0.15 | (0.00, 0.51) | 0.14 | (0.00, 0.42) |
| Monandrous species | 32 | 16 | 0.22 | (-0.02, 0.45) | 0.067 | 0.24 | (0.00, 0.67) | 0.53 | (0.01, 0.91) | 0.12 | (0.00, 0.47) |
| Polyandrous species | 79 | 56 | 0.40 | (0.11, 0.67) | 0.010 | 0.72 | (0.40, 0.95) | 0.09 | (0.00, 0.31) | 0.10 | (0.00, 0.27) |

**Table 2.**
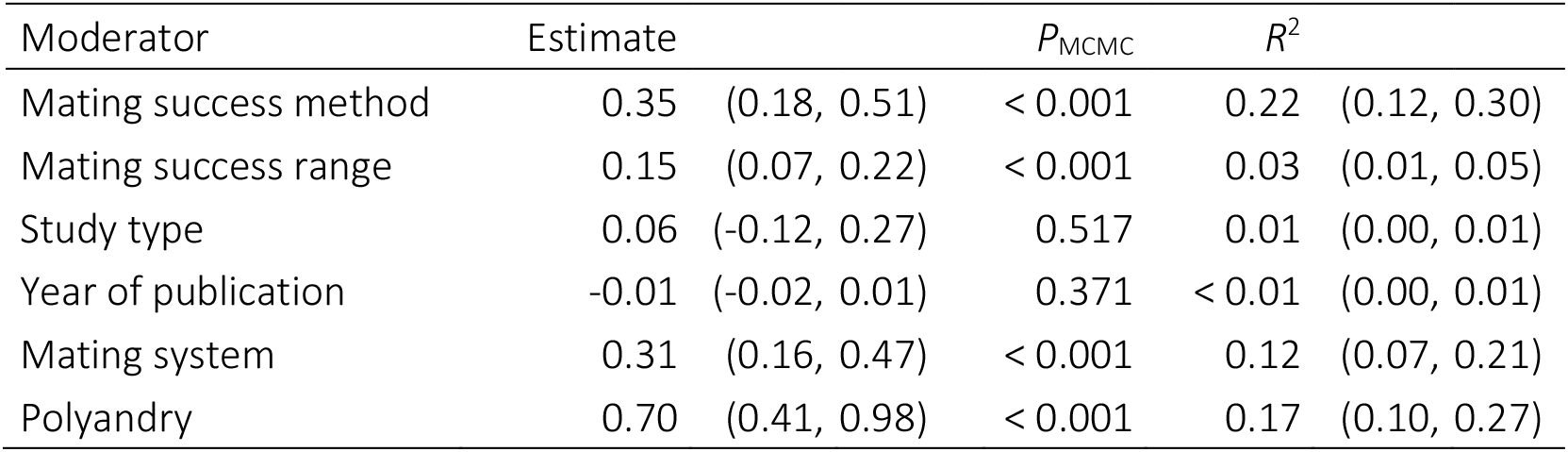
Predictors of inter-specific variation in female Bateman gradients. Methodological moderators include mating success method (copulatory *versus* genetic mating success), mating success range (including *versus* excluding zero mating success category), study type (field *versus* lab) and year of publication (continuous variable). Effect of mating system contrasts polyandrous and monandrous species. Effect of polyandry estimates the relationship between the female Bateman gradient and the proportion of polyandrous females in the population. Model estimates (i.e., estimated difference between groups) are shown as posterior modes with 95% Highest Posterior Density (HPD) intervals obtained from phylogenetically controlled General Linear-Mixed Effects Models. The variance explained by the moderator variable is given as the marginal *R*^2^ with 95% HPD intervals.

Importantly, estimates of the strength of sexual selection differed between monandrous and polyandrous species. As predicted, polyandrous species show stronger Bateman gradients compared to monandrous species (Table 2 & S2; Figure 1B). This effect is also supported by an alternative analysis indicating a positive relationship between the actual level of polyandry and the Bateman gradient (Table 2 & S2).

We observed a statistically significant signal for funnel plot asymmetry (rank correlation test: Kendall’s tau = −0.146, *P* = 0.023), partly driven by an underrepresentation of low-powered studies with high effect sizes (Figure S6). This may indicate a publication bias towards small studies with weak or non-significant female Bateman gradients. However, a clear-cut interpretation of funnel plot asymmetry is difficult especially in the presence of unexplained heterogeneity. Finally, we did not detect an effect of publication year suggesting the absence of the so-called bandwagon effect (Table 2 and S2).

## Discussion

Darwin’s male-centred perspective predominates our general perception of sexual reproduction. In support of the so-called Darwin-Bateman paradigm (*28*, *44*), sexual selection has indeed been found to act typically stronger in males compared to females (*6*), but the significance of the sex-role concept emphasizing pervasive sex differences in mate competition and mate choice is still subject to an ongoing controversy across disciplines (*14*, *28*, *45*–*48*). Our study aims at illuminating this debate by providing quantitative evidence that sexual selection on females is the norm rather than the exception across the animal tree of life. Specifically, our results document that females – just as widely assumed for males – typically benefit from having more than one mating partner (indicated by a positive Bateman gradient). As a consequence, selection is also expected to favour the evolution of female traits that promote the acquisition of mating partners. However, given the previously documented higher benefit of mating in males (*6*), sexual selection on females may often operate more hiddenly, leading to the evolution of less conspicuous ornaments and armaments compared to males.

Importantly, our results are robust with respect to different methodological approaches used to estimate the Bateman gradient. Even after exclusion of study designs that are prone to overestimate the relationship between mating and reproductive success (i.e., those using genetic parentage rather than behavioural observations to quantify mating success; (*27*)), we detected an overall positive Bateman gradient. Moreover, our findings suggest that the positive relationship between mating success and reproductive success in females is not only driven by the benefit of having at least a single mating – which is arguably rather trivial – but also by the benefit of having an additional mating. Hence, despite multifaceted evidence that mating can incur costs for females (*49*, *50*) and observations that females can be sperm limited (*51*), reproductive success may often be maximized at a mating rate that is higher than required for fertilising all eggs. Remarkably, we also detected a correlation between the Bateman gradient and the mating system, implying that species showing a steeper female Bateman gradient tend to be more polyandrous. Even if our comparative approach does not allow inference of causality, this finding highly suggests that directional positive selection on mating success is effective and translates into higher mating rates in females as predicted by sexual selection theory (*20*, *30*).

The overall positive effect of mating on reproductive success in females raises the question of what the underlying benefits of an additional mating actually are. A previous meta-analysis on the evolution of polyandry in insects found that fitness benefits of multiple mating in females are especially high in species in which males provide nuptial feeding (*52*). However, females may benefit from mating in manifold ways including other direct benefits such as paternal care or indirectly in terms of genetic benefits (*53*, *54*). The relative importance of these diverse benefits and their role in promoting sexual selection on females still remain to be explored at a broader macro-evolutionary scale.

Our study relies on the premise that the Bateman gradient provides a meaningful quantitative proxy for the strength of sexual selection. While there is compelling theoretical and empirical support for this assertion, especially in the context of interspecific comparisons (*27*, *28*, *34*, *55*), it also comes with limitations. First, the Bateman gradient captures primarily pre-copulatory episodes of sexual selection. While there is a clear scope for egg competition and post-copulatory “cryptic” male choice (*17*), internal fertilisation in females is emblematic for the vast majority of terrestrial animals. This is likely to limit post-copulatory competition among females, which makes the female Bateman gradient a less incomplete proxy for the total strength of sexual selection compared to males. Second, the Bateman gradient captures only the upper potential of actual phenotypic selection (*27*), therefore, our study cannot provide a trait-based perspective on female sexual selection. However, our results suggest that the potential for the evolution of sexually selected traits in females is more widespread than previously thought, which challenges arguments that ornamentation in females evolves mainly as a by-product of sexual selection on males (*56*). The quantity of all documented examples of sexually selected traits in females may reflect an imbalance in research efforts and possibly represents just the tip of the iceberg of what can be found in nature.

Collectively, our study contributes to a more nuanced view on sexual selection and sex differences in general. Darwinian sex roles may predominate the animal tree of life in the sense that sexual selection is typically stronger on males compared to females (*6*) but our meta-analysis questions the view that females are typically coy and passive. Sexual selection on females should not be considered a rare phenomenon but instead be acknowledged as widespread across animals.

## Acknowledgements

We are very grateful to the authors of all primary studies for making their research accessible, especially those providing additional data allowing us to compute Bateman gradients. We also thank Julie Collet and Karoline Fritzsche for providing unpublished data to quantify the level of polyandry of their study organisms.

## Author contributions

TJ and LMO conceived the study. SF, LW, LMO and TJ designed the meta-analysis. SF and TJ collected the data with help from LW and LMO. SF and TJ performed the statistical analyses. SF and TJ wrote the paper with the help from LW and LMO.

## Data and materials availability

After acceptance of the article, all data will be uploaded on the Dryad digital repository (https://v1.datadryad.org/).

