## Supplementary Figures and Tables for "Sexual selection in females across the animal tree of life"

**Contents**

Figure S1 – S6. (pp. 2 – 7)

Table S1 – S3 (pp. 8 – 15)

Supplementary Data: List of primary studies (pp. 16– 20)

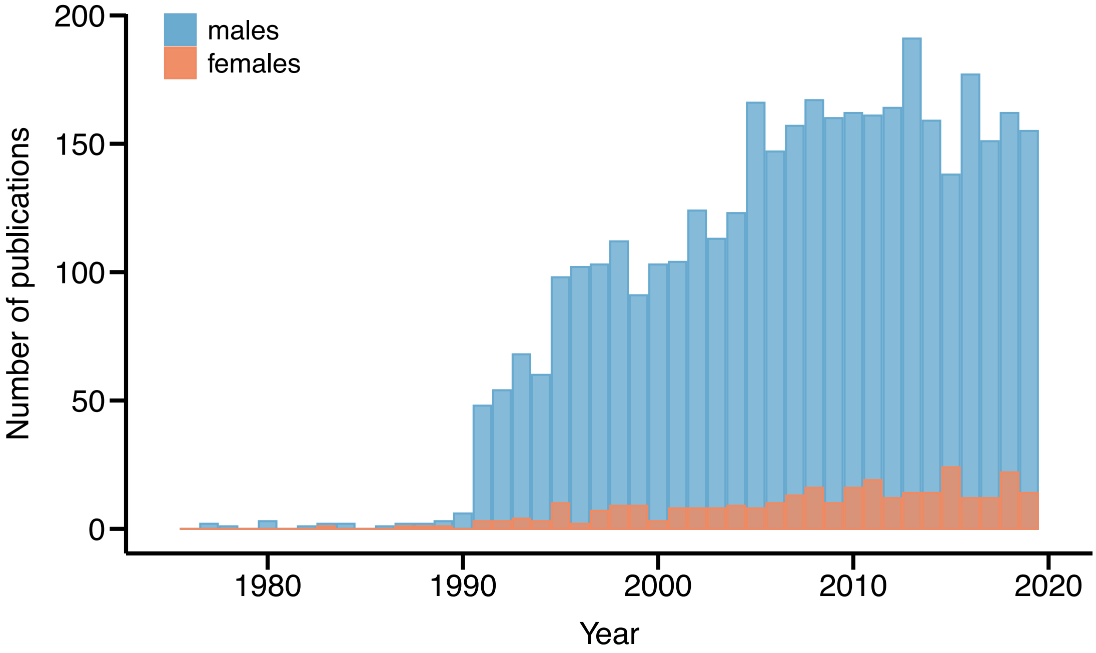


**Figure S1. Imbalance between studies of sexual selection in males and females.** Bars indicate a strong male bias in the number of published articles on sexual selection indexed in ISI Web of Science (Clarivate Analytics) between 1900-2020. Data obtained from topic search using the search terms "sexual selection AND (male choice OR female competition)" for female and "sexual selection AND female choice OR male competition)" for male search. This is not meant to provide an exhaustive search of publications on sexual selection but to showcase the publication bias towards male studies focusing on Darwinian sexual selection in terms of competition for and choice of mating partners.

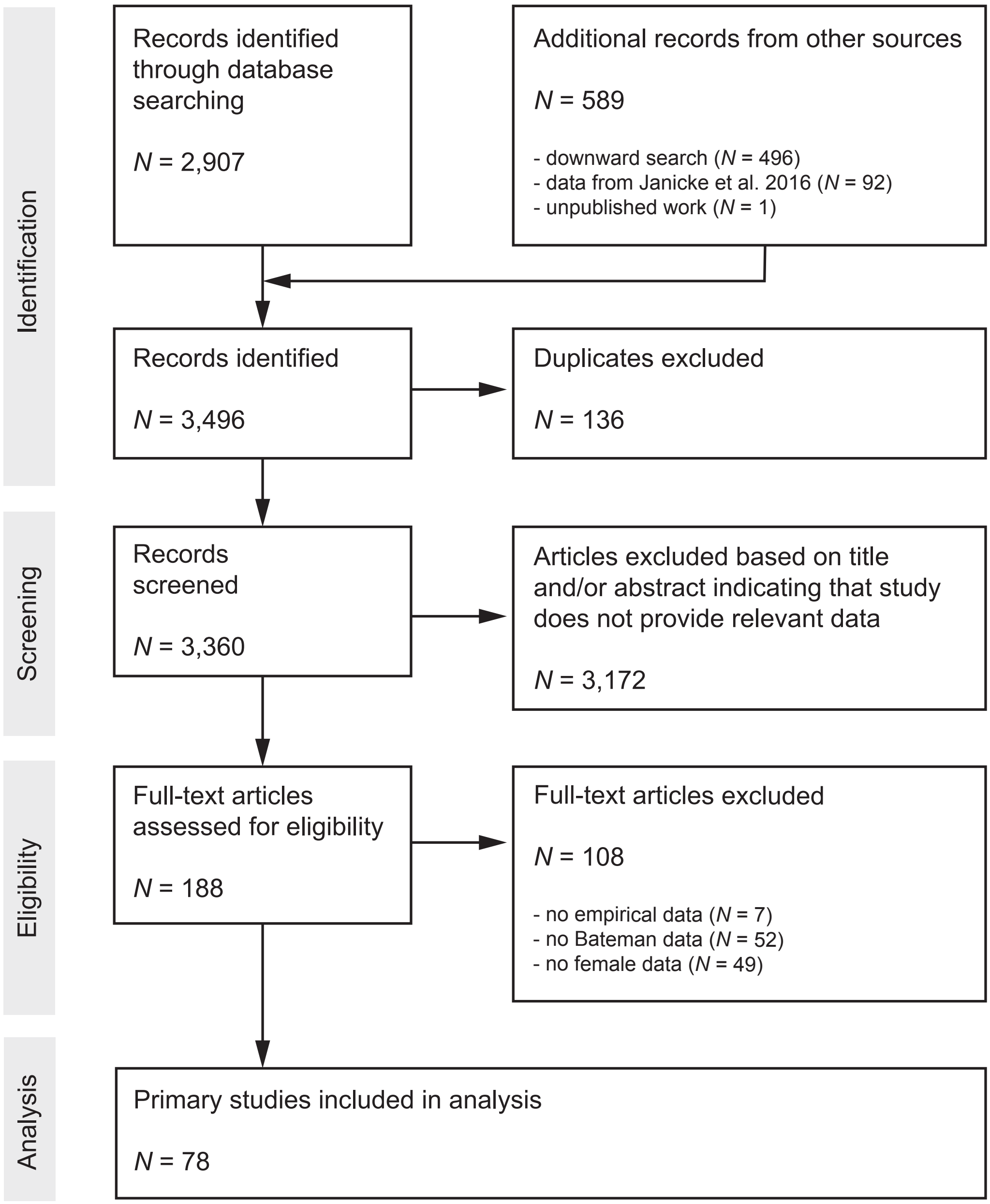


**Figure S2. Preferred Reporting Items for Systematic Reviews and Meta-Analyses (PRISMA) Diagram.** Flow chart maps the number of records identified during the different phases of the systematic literature search.

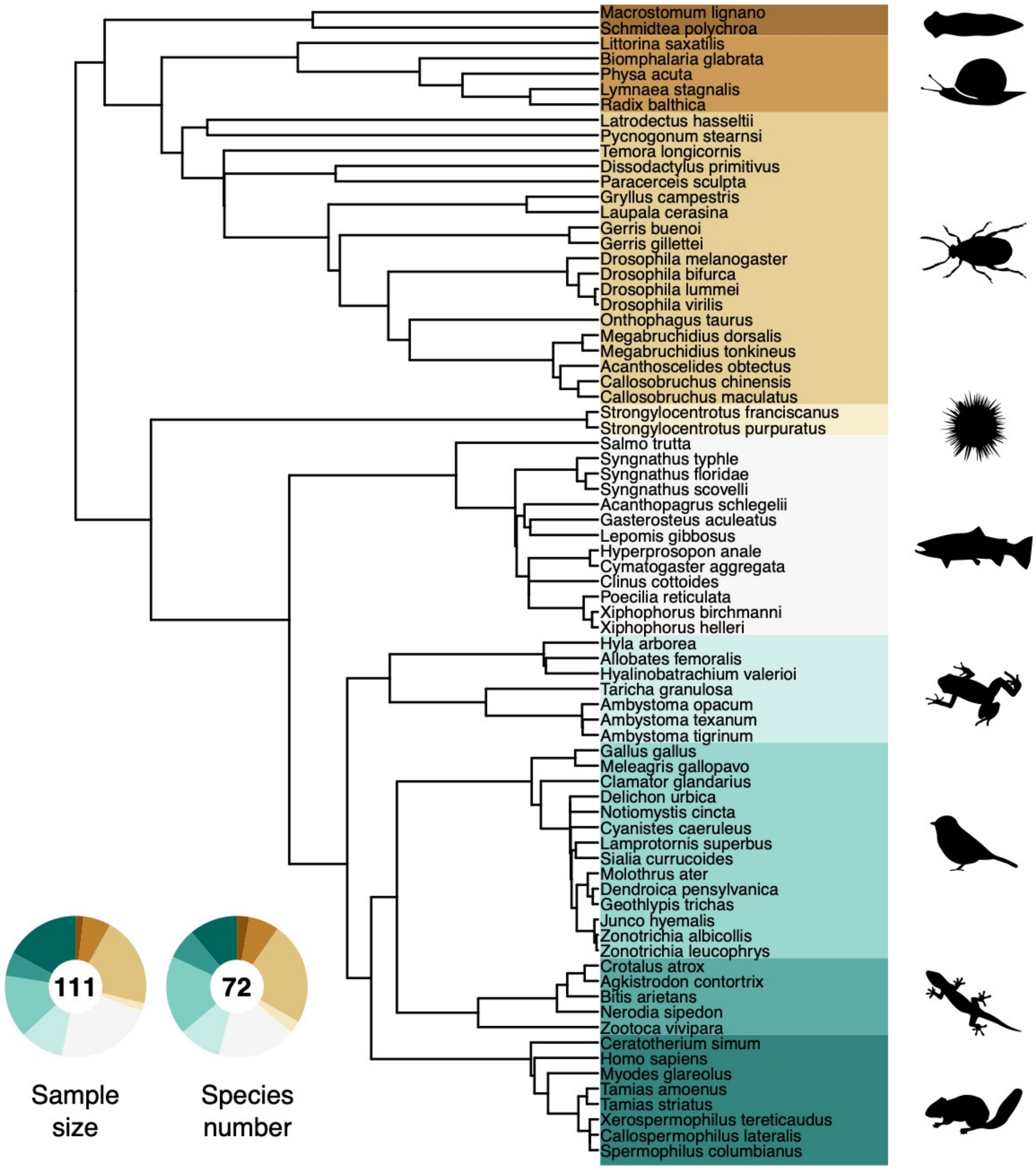


**Figure S3. Phylogenetic tree of all sampled species.** Doughnut charts show the relative fraction of the sampled effect sizes (i.e., number of Bateman gradients) and the number of species.

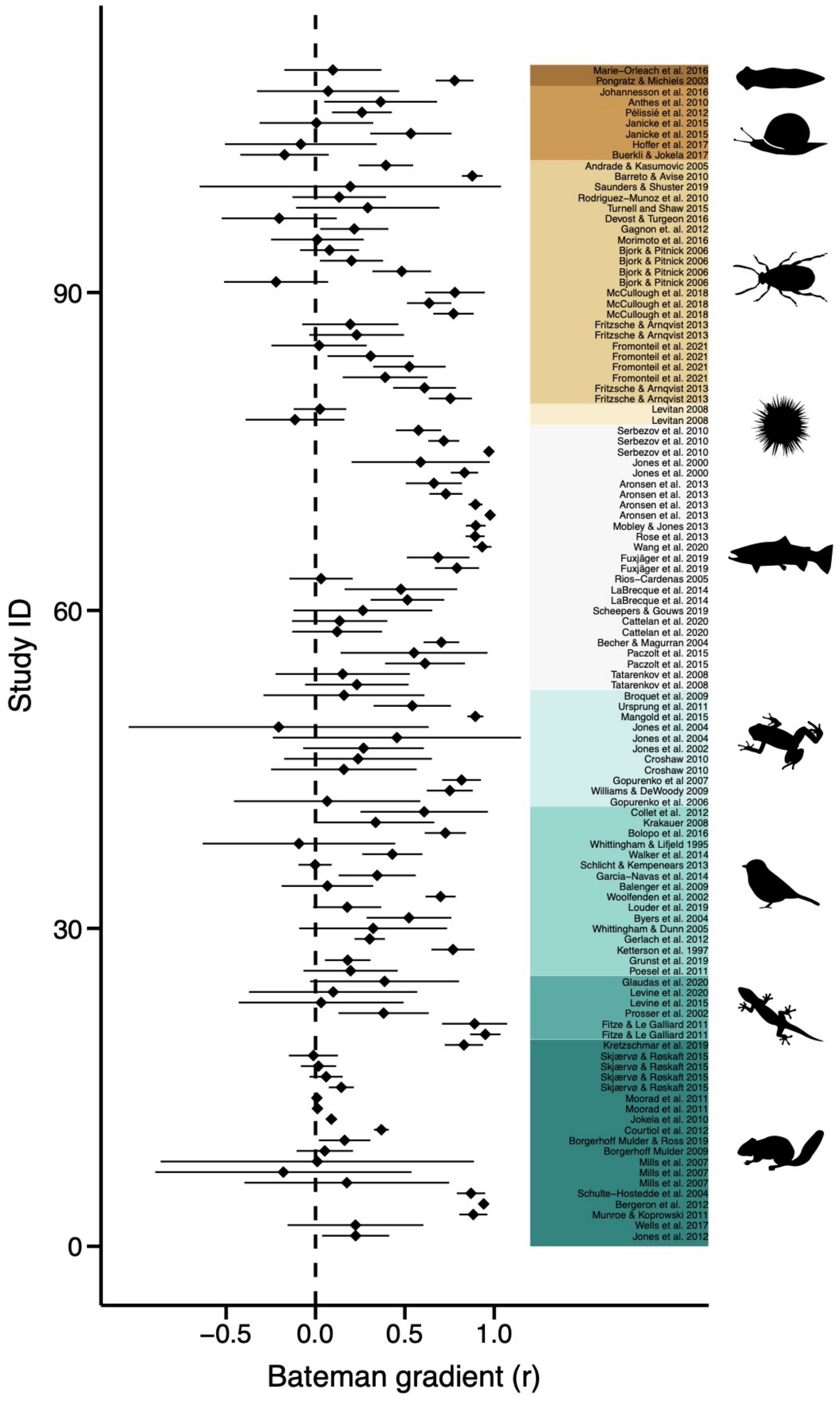


**Figure S4. Forest plot of all sampled effect sizes**. Effect sizes (Pearson correlation coefficient of Bateman gradients) with 95% confidence limits are shown in phylogenetic order.

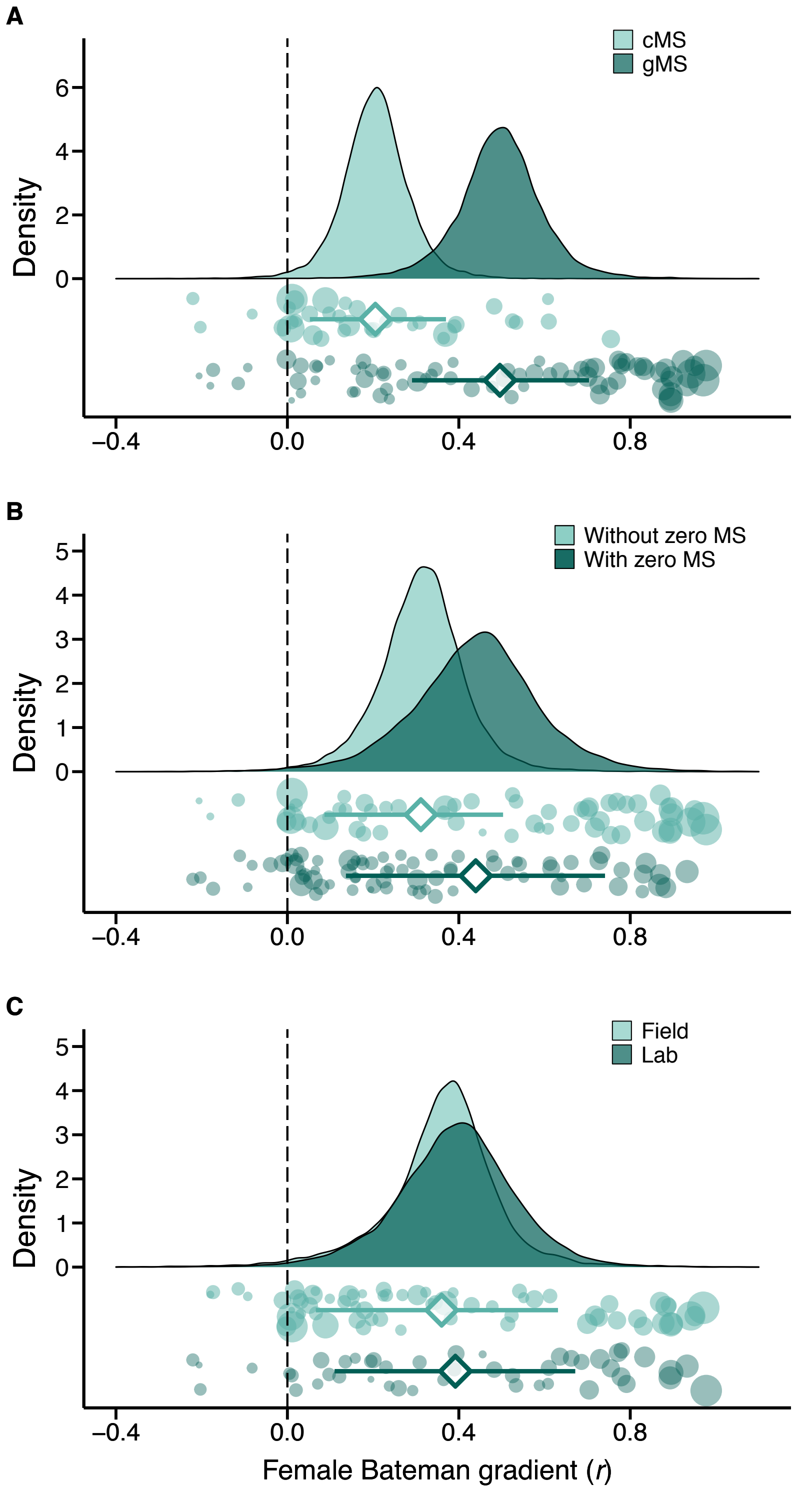


**Figure S5. Methodological predictors of female Bateman gradients.** Raincloud charts showing effects of mating success method (*cMS*: copulatory mating success, *gMS*: genetic mating success), mating success range (with or without zero mating success (*MS*) category) and study type (field *versus* laboratory studies) on female Bateman gradients (for statistical analysis see Table 2 and S2).

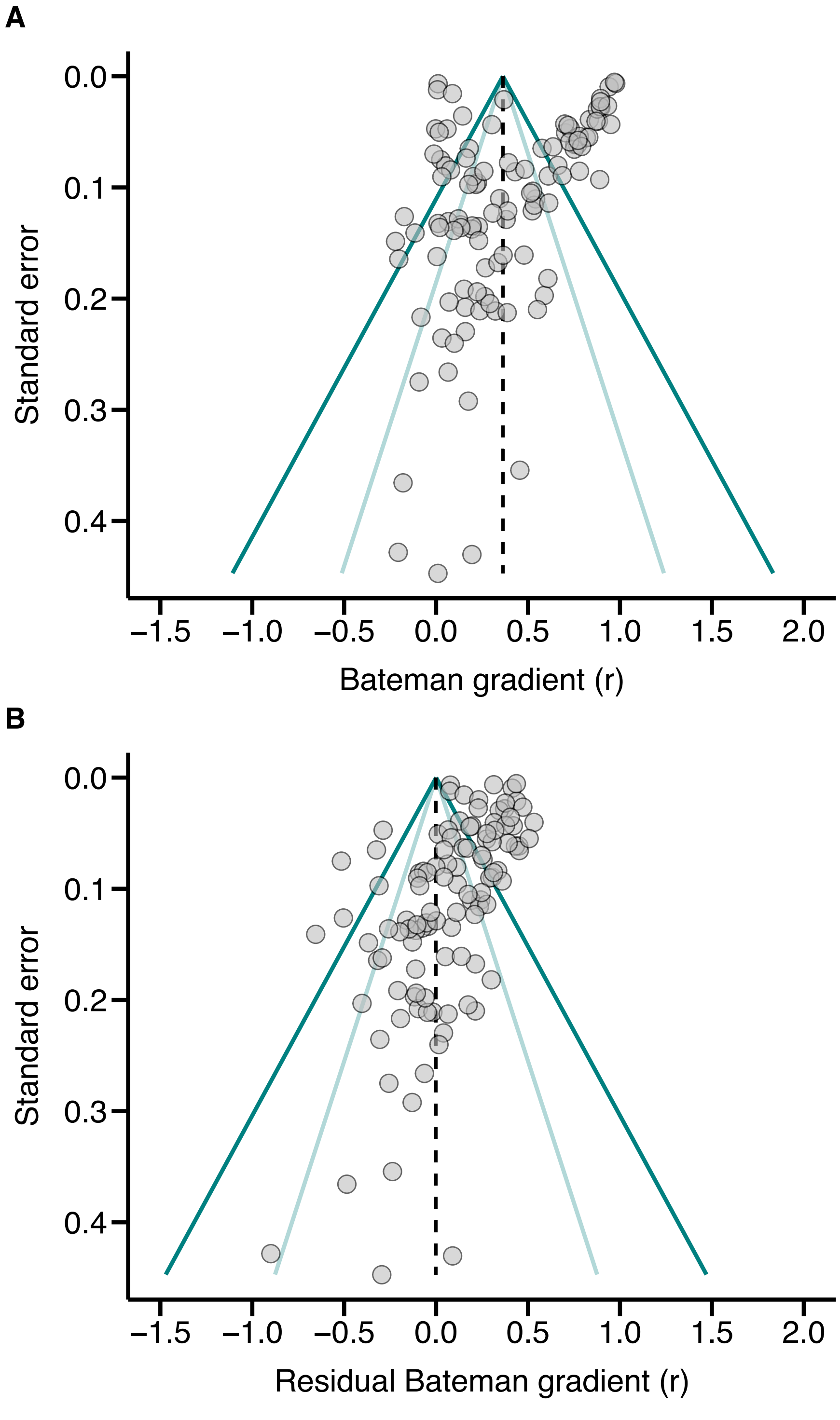


**Figure S6. Funnel plots.** Data are shown for raw values (A) and for meta-analytic residuals obtained from multivariate linear mixed-effects models accounting for all tested moderator variables. Dashed lines indicate the estimated global effect size. Dark and light blue solid lines denote the expected 95% and 99% confidence limits purely due to sampling heterogeneity. Asymmetries along the global effect size may reflect publication biases.

**Table S1. Global tests of sexual selection in females using the restricted maximum likelihood (REML) approach.** Results of intercept-only phylogenetically controlled General Linear-Mixed Effects Models are shown for the entire dataset (global model) and subsets with respect to mating success method (copulatory versus genetic), mating success range (including versus excluding zero mating success category), study type (laboratory versus field studies) and mating system (monandrous versus polygamous species). Table shows number of effect sizes (*k*), number of species (*N*) and estimates of *r* together with 95% confidence intervals (in brackets).

| Model | *k* | $N_{Species}$ | Global effect size *r* | | *z*-value | *P*-value |
|---|---|---|---|---|---|---|
| Global model | 111 | 72 | 0.36 | (0.10, 0.63) | 2.662 | 0.008 |
| Copulatory mating success | 37 | 21 | 0.21 | (0.06, 0.35) | 2.854 | 0.004 |
| Genetic mating success | 76 | 54 | 0.50 | (0.31, 0.68) | 5.301 | < 0.001 |
| Including zero mating success | 61 | 37 | 0.44 | (0.14, 0.73) | 2.905 | 0.004 |
| Excluding zero mating success | 73 | 55 | 0.30 | (0.13, 0.48) | 3.411 | 0.001 |
| Laboratory studies | 46 | 28 | 0.39 | (0.11, 0.66) | 2.766 | 0.006 |
| Field studies | 65 | 45 | 0.39 | (0.30, 0.48) | 8.324 | < 0.001 |
| Monandrous species | 32 | 16 | 0.23 | (0.09, 0.37) | 3.162 | 0.002 |
| Polyandrous species | 79 | 56 | 0.40 | (0.11, 0.69) | 2.727 | 0.006 |

**Table S2. Predictors of inter-specific variation in female Bateman gradients using restricted maximum likelihood (REML) approach.** Methodological moderators include mating success method (copulatory *versus* genetic mating success), mating success range (including *versus* excluding mating success category), study type (field *versus* lab) and year of publication (continuous variable). Effect of mating system contrasts polyandrous and monandrous species. Effect of polyandry (continuous variable) estimates the relationship between the female Bateman gradient and the proportion of polyandrous females in the population. Phylogenetically controlled multilevel meta-analytic single predictor models are shown with Omnibus tests (Wald-type chi-square test) and McFadden's $R^2$.

| Moderator | Estimate ± SE | $Q_M$ | *P*-value | $R^2$ |
|---|---|---|---|---|
| Mating success method | **0.30 ± 0.09** | 10.84 | 0.001 | 0.20 |
| Mating success range | **0.18 ± 0.06** | 7.65 | 0.006 | 0.15 |
| Study type | 0.07 ± 0.10 | 0.51 | 0.475 | 0.02 |
| Year | -0.01 ± 0.01 | 0.84 | 0.358 | 0.01 |
| Mating system | **0.31 ± 0.08** | 16.35 | < 0.001 | 0.32 |
| Polyandry | **0.68 ± 0.14** | 23.25 | < 0.001 | 0.46 |

Supplementary Data

### Mating system classification

**Table S3.** Estimates of polyandry and mating system classification (monandrous *versus* polyandrous) of the 72 sampled species in alphabetical order.

| Species | Polyandry | Mating system | Reference |
|---|---|---|---|
| *Acanthopagrus schlegelii* | 0.920 | Polyandrous | (Wang *et al.* 2020) |
| *Acanthoscelides obtectus* | 0.551 | Polyandrous | Fromonteil et al. (in prep.)[1] |
| *Agkistrodon contortrix* | 0.520 | Polyandrous | (Levine *et al.* 2015) |
| *Allobates femoralis* | 0.571 | Polyandrous | (Ursprung *et al.* 2011) |
| *Ambystoma opacum* | 0.294 | Monandrous | (Croshaw 2010) |
| *Ambystoma texanum* | 0.857 | Polyandrous | (Gopurenko *et al.* 2007) |
| *Ambystoma tigrinum* | 0.467 | Monandrous | (Gopurenko *et al.* 2006) |
| *Biomphalaria glabrata* | 0.654 | Polyandrous | (Anthes *et al.* 2010) |
| *Bitis arietans* | 0.941 | Polyandrous | (Glaudas *et al.* 2020) |
| *Callosobruchus chinensis* | 0.680 | Polyandrous | (Fritzsche & Arnqvist 2013)[2] |
| *Callosobruchus maculatus* | 1.000 | Polyandrous | (Fritzsche & Arnqvist 2013)[2] |
| *Callospermophilus lateralis* | 0.630 | Polyandrous | (Wells *et al.* 2017) |
| *Ceratotherium simum* | 0.697 | Polyandrous | (Kretzschmar *et al.* 2020) |
| *Clamator glandarius* | 0.308 | Monandrous | (Bolopo *et al.* 2017) |
| *Clinus cottoides* | 0.826 | Polyandrous | (Scheepers & Gouws 2019) |
| *Crotalus atrox* | 0.400 | Monandrous | (Levine *et al.* 2020) |
| *Cyanistes caeruleus* | 0.470 | Monandrous | (Schlicht & Kempenaers 2013) |
| *Cymatogaster aggregata* | 0.800 | Polyandrous | (LaBrecque *et al.* 2014) |
| *Delichon urbica* | 0.235 | Monandrous | (Whittingham & Lifjeld 1995) |
| *Dendroica pensylvanica* | 0.606 | Polyandrous | (Byers *et al.* 2004) |
| *Drosophila bifurca* | 0.326 | Monandrous | (Mery & Joly 2002)[3] |
| *Drosophila lummei* | 0.753 | Polyandrous | (Bjork & Pitnick 2006)[3] |
| *Drosophila melanogaster* | 0.759 | Polyandrous | (Morimoto *et al.* 2016) |
| *Drosophila virilis* | 0.753 | Polyandrous | (Bjork & Pitnick 2006)[3] |
| *Gallus gallus* | 0.950 | Polyandrous | (Collet *et al.* 2012)[2] |
| *Gasterosteus aculeatus* | 0.783 | Polyandrous | (Fuxjager *et al.* 2019) |
| *Geothlypis trichas* | 0.654 | Polyandrous | (Whittingham & Dunn 2005) |
| *Gerris buenoi* | 1.000 | Polyandrous | (Devost & Turgeon 2016) |
| *Gerris gillettei* | 0.776 | Polyandrous | (Gagnon *et al.* 2012) |
| *Gryllus campestris* | 0.809 | Polyandrous | (Rost & Honegger 1987) |
| *Homo sapiens* | 0.012 | Monandrous | (Larmuseau *et al.* 2016) |
| *Hyalinobatrachium valerioi* | 0.736 | Polyandrous | (Mangold *et al.* 2015) |
| *Hyla arborea* | 0.158 | Monandrous | (Paczolt *et al.* 2015) |
| *Hyperprosopon anale* | 1.000 | Polyandrous | (LaBrecque *et al.* 2014) |
| *Junco hyemalis* | 0.439 | Monandrous | (Ketterson *et al.* 1997) |

| Species | Polyandry | Mating system | Reference |
|---|---|---|---|
| *Latrodectus hasselti* | 0.667 | Polyandrous | (Andrade & Kasumovic 2005) |
| *Laupala cerasina* | 1.000 | Polyandrous | (Turnell & Shaw 2015) |
| *Lepomis gibbosus* | 0.244 | Monandrous | (Rios-Cardenas 2005) |
| *Littorina saxatilis* | 0.680 | Polyandrous | (Johannesson *et al.* 2016) |
| *Lymnaea stagnalis* | 0.588 | Polyandrous | (Nakadera *et al.* 2017) |
| *Macrostomum lignano* | 1.000 | Polyandrous | (Marie-Orleach *et al.* 2016) |
| *Megabruchidius dorsalis* | 1.000 | Polyandrous | (Fritzsche & Arnqvist 2013)[2] |
| *Megabruchidius tonkineus* | 0.977 | Polyandrous | (Fritzsche & Arnqvist 2013)[2] |
| *Meleagris gallopavo* | 0.450 | Monandrous | (Krakauer 2008) |
| *Molothrus ater* | 0.455 | Monandrous | (Strausberger & Ashley 2003) |
| *Myodes glareolus* | 0.353 | Monandrous | (Ratkiewicz & Borkowska 2000) |
| *Nerodia sipedon* | 0.556 | Polyandrous | (Prosser *et al.* 2002) |
| *Notiomystis cincta* | 0.836 | Polyandrous | (Walker *et al.* 2014) |
| *Onthophagus taurus* | 0.789 | Polyandrous | (McCullough *et al.* 2018) |
| *Paracerceis sculpta* | 0.667 | Polyandrous | (Saunders & Shuster 2019) |
| *Physa acuta* | 0.789 | Polyandrous | (Pelissie *et al.* 2012) |
| *Poecilia reticulata* | 0.710 | Polyandrous | (Cattelan *et al.* 2020) |
| *Pycnogonum stearnsi* | 0.686 | Polyandrous | (Barreto & Avise 2010) |
| *Radix balthica* | 0.500 | Polyandrous | (Burkli & Jokela 2017) |
| *Salmo trutta* | 0.680 | Polyandrous | (Largiader *et al.* 2001) |
| *Schmidtea polychroa* | 0.918 | Polyandrous | (Pongratz & Michiels 2003) |
| *Sialia currucoides* | 0.342 | Monandrous | (Brouwer & Griffith 2019) |
| *Spermophilus columbianus* | 0.343 | Monandrous | (Jones *et al.* 2012) |
| *Strongylocentrotus franciscanus* | 0.990 | Polyandrous | (Levitan 2008) |
| *Strongylocentrotus purpuratus* | 0.990 | Polyandrous | (Levitan 2008) |
| *Syngnathus floridae* | 0.633 | Polyandrous | (Jones & Avise 1997) |
| *Syngnathus scovelli* | 0.615 | Polyandrous | (Jones *et al.* 2001) |
| *Syngnathus typhle* | 0.528 | Polyandrous | (Jones *et al.* 2000) |
| *Tamias amoenus* | 0.595 | Polyandrous | (Schulte-Hostedde *et al.* 2004) |
| *Tamias striatus* | 0.650 | Polyandrous | (Bergeron *et al.* 2012) |
| *Taricha granulosa* | 0.548 | Polyandrous | (Jones *et al.* 2004) |
| *Xerospermophilus tereticaudus* | 0.935 | Polyandrous | (Munroe & Koprowski 2011) |
| *Xiphophorus birchmanni* | 0.839 | Polyandrous | (Paczolt *et al.* 2015) |
| *Xiphophorus helleri* | 0.638 | Polyandrous | (Tatarenkov *et al.* 2008) |
| *Zonotrichia albicollis* | 0.306 | Monandrous | (Grunst *et al.* 2019) |
| *Zonotrichia leucophrys* | 0.346 | Monandrous | (Poesel *et al.* 2011) |
| *Zootoca vivipara* | 0.633 | Polyandrous | (Laloi *et al.* 2004) |

[1] Unpublished data: S. Fromonteil.
[2] Authors of primary studies kindly provided on request attentional data to compute polyandry.
[3] References could only be found for classification of the mating system. Polyandry was extrapolated from the average level of polyandry observed in polygynous or monandrous species, respectively.

#### References – Mating system classification


Andrade, M.C.B. & Kasumovic, M.M. (2005). Terminal investment strategies and male mate choice: Extreme tests of Bateman. *Integrative and Comparative Biology*, 45, 838-847.

Anthes, N., David, P., Auld, J.R., Hoffer, J.N., Jarne, P., Koene, J.M. *et al.* (2010). Bateman gradients in hermaphrodites: an extended approach to quantify sexual selection. *American Naturalist*, 176, 249-263.

Barreto, F.S. & Avise, J.C. (2010). Quantitative measures of sexual selection reveal no evidence for sex-role reversal in a sea spider with prolonged paternal care. *Proceedings of the Royal Society B-Biological Sciences*, 277, 2951-2956.

Bergeron, P., Montiglio, P.O., Reale, D., Humphries, M.M. & Garant, D. (2012). Bateman gradients in a promiscuous mating system. *Behav. Ecol. Sociobiol.*, 66, 1125-1130.

Bjork, A. & Pitnick, S. (2006). Intensity of sexual selection along the anisogamy-isogamy continuum. *Nature*, 441, 742-745.

Bolopo, D., Canestrari, D., Martinez, J.G., Roldan, M., Macias-Sanchez, E., Vila, M. *et al.* (2017). Flexible mating patterns in an obligate brood parasite. *Ibis*, 159, 103-112.

Brouwer, L. & Griffith, S.C. (2019). Extra-pair paternity in birds. *Mol. Ecol.*, 28, 4864-4882.

Burkli, A. & Jokela, J. (2017). Increase in multiple paternity across the reproductive lifespan in a sperm-storing, hermaphroditic freshwater snail. *Mol. Ecol.*, 26, 5264-5278.

Byers, B.E., Mays, H.L., Stewart, I.R.K. & Westneat, D.F. (2004). Extrapair paternity increases variability in male reproductive success in the chestnut-slded warbler (*Dendroica pensylvanica*), a socially monogamous songbird. *Auk*, 121, 788-795.

Cattelan, S., Evans, J.P., Garcia-Gonzalez, F., Morbiato, E. & Pilastro, A. (2020). Dietary stress increases the total opportunity for sexual selection and modifies selection on condition-dependent traits. *Ecol. Lett.*, 23, 447-456.

Collet, J., Richardson, D.S., Worley, K. & Pizzari, T. (2012). Sexual selection and the differential effect of polyandry. *Proc. Natl. Acad. Sci. U. S. A.*, 109, 8641-8645.

Croshaw, D.A. (2010). Quantifying sexual selection: a comparison of competing indices with mating system data from a terrestrially breeding salamander. *Biological Journal of the Linnean Society*, 99, 73-83.

Devost, E. & Turgeon, J. (2016). The combined effects of pre- and post-copulatory processes are masking sexual conflict over mating rate in *Gerris buenoi*. *Journal of Evolutionary Biology*, 29, 167-177.

Fritzsche, K. & Arnqvist, G. (2013). Homage to Bateman: sex roles predict sex differences in sexual selection. *Evolution; international journal of organic evolution*, 67, 1926-1936.

Fuxjager, L., Wanzenbock, S., Ringler, E., Wegner, K.M., Ahnelt, H. & Shama, L.N.S. (2019). Within-generation and transgenerational plasticity of mate choice in oceanic stickleback under climate change. *Philos. Trans. R. Soc. B-Biol. Sci.*, 374, 12.

Gagnon, M.-C., Duchesne, P. & Turgeon, J. (2012). Sexual conflict in *Gerris gillettei* (Insecta: Hemiptera): influence of effective mating rate and morphology on reproductive success. *Canadian Journal of Zoology*, 90, 1297-1306.

Glaudas, X., Rice, S.E., Clark, R.W. & Alexander, G.J. (2020). The intensity of sexual selection, body size and reproductive success in a mating system with male-male combat: is bigger better? *Oikos*, 129, 998-1011.

Gopurenko, D., Williams, R.N. & DeWoody, J.A. (2007). Reproductive and mating success in the small-mouthed salamander (*Ambystoma texanum*) estimated via microsatellite parentage analysis. *Evolutionary Biology*, 34, 130-139.

Gopurenko, D., Williams, R.N., McCormick, C.R. & DeWoody, J.A. (2006). Insights into the mating habits of the tiger salamander (*Ambystoma tigrinum tigrinum*) as revealed by genetic parentage analyses. *Mol. Ecol.*, 15, 1917-1928.

Grunst, A.S., Grunst, M.L., Korody, M.L., Forrette, L.M., Gonser, R.A. & Tuttle, E.M. (2019). Extrapair mating and the strength of sexual selection: insights from a polymorphic species. *Behav. Ecol.*, 30, 278-290.

Johannesson, K., Saltin, S.H., Charrier, G., Ring, A.K., Kvarnemo, C., Andre, C. *et al.* (2016). Non-random paternity of offspring in a highly promiscuous marine snail suggests postcopulatory sexual selection. *Behav. Ecol. Sociobiol.*, 70, 1357-1366.

Jones, A.G., Arguello, J.R. & Arnold, S.J. (2004). Molecular parentage analysis in experimental newt populations: The response of mating system measures to variation in the operational sex ratio. *American Naturalist*, 164, 444-456.

Jones, A.G. & Avise, J.C. (1997). Polygynandry in the dusky pipefish *Syngnathus floridae* revealed by microsatellite DNA markers. *Evolution; international journal of organic evolution*, 51, 1611-1622.

Jones, A.G., Rosenqvist, G., Berglund, A. & Avise, J.C. (2000). Mate quality influences multiple maternity in the sex-role-reversed pipefish *Syngnathus typhle*. *Oikos*, 90, 321-326.

Jones, A.G., Walker, D. & Avise, J.C. (2001). Genetic evidence for extreme polyandry and extraordinary sex-role reversal in a pipefish. *Proceedings. Biological sciences / The Royal Society*, 268, 2531-2535.

Jones, P.H., Van Zant, J.L. & Dobson, F.S. (2012). Variation in reproductive success of male and female Columbian ground squirrels (*Urocitellus columbianus*). *Can. J. Zool.-Rev. Can. Zool.*, 90, 736-743.

Ketterson, E.D., Parker, P.G., Raouf, S.A., Nolan Jr, V., Ziegenfus, C. & Chandler, C.H. (1997). The relative impact of extra-pair fertilizations on variation in male and female reproductive success in dark-eyed juncos (*Junco hyemaus*). In: *Avian Reproductive Tactics: Female and Male Perspectives* (eds. Parker, PG & Burley, NT), pp. 81-101.

Krakauer, A.H. (2008). Sexual selection and the genetic mating system of Wild Turkeys. *Condor*, 110, 1-12.

Kretzschmar, P., Auld, H., Boag, P., Ganslosser, U., Scott, C., de Groot, P.J.V. *et al.* (2020). Mate choice, reproductive success and inbreeding in white rhinoceros: New insights for conservation management. *Evol. Appl.*, 13, 699-714.

LaBrecque, J.R., Alva-Campbell, Y.R., Archambeault, S. & Crow, K.D. (2014). Multiple paternity is a shared reproductive strategy in the live-bearing surfperches (Embiotocidae) that may be associated with female fitness. *Ecol. Evol.*, 4, 2316-2329.

Laloi, D., Richard, M., Lecomte, J., Massot, M. & Clobert, J. (2004). Multiple paternity in clutches of common lizard *Lacerta vivipara*: data from microsatellite markers. *Mol. Ecol.*, 13, 719-723.

Largiader, C.R., Estoup, A., Lecerf, F., Champigneulle, A. & Guyomard, R. (2001). Microsatellite analysis of polyandry and spawning site competition in brown trout (*Salmo trutta* L.). *Genetics Selection Evolution*, 33, S205-S222.

Larmuseau, M.H.D., Matthijs, K. & Wenseleers, T. (2016). Cuckolded fathers rare in human populations. *Trends in Ecology & Evolution*, 31, 327-329.

Levine, B.A., Schuett, G.W., Clark, R.W., Repp, R.A., Herrmann, H.W. & Booth, W. (2020). No evidence of male-biased sexual selection in a snake with conventional Darwinian sex roles. *R. Soc. Open Sci.*, 7, 10.

Levine, B.A., Smith, C.F., Schuett, G.W., Douglas, M.R., Davis, M.A. & Douglas, M.E. (2015). Bateman-Trivers in the 21st Century: sexual selection in a North American pitviper. *Biological Journal of the Linnean Society*, 114, 436-445.

Levitan, D.R. (2008). Gamete traits influence the variance in reproductive success, the intensity of sexual selection, and the outcome of sexual conflict among congeneric sea urchins. *Evolution; international journal of organic evolution*, 62, 1305-1316.

Mangold, A., Trenkwalder, K., Ringler, M., Hoedl, W. & Ringler, E. (2015). Low reproductive skew despite high male-biased operational sex ratio in a glass frog with paternal care. *Bmc Evolutionary Biology*, 15.

Marie-Orleach, L., Janicke, T., Vizoso, D.B., David, P. & Scharer, L. (2016). Quantifying episodes of sexual selection: Insights from a transparent worm with fluorescent sperm. *Evolution; international journal of organic evolution*, 70, 314-328.

McCullough, E.L., Buzatto, B.A. & Simmons, L.W. (2018). Population density mediates the interaction between pre- and postmating sexual selection. *Evolution; international journal of organic evolution*, 72, 893-905.

Mery, F. & Joly, D. (2002). Multiple mating, sperm transfer and oviposition pattern in the giant sperm species, *Drosophila bifurca*. *Journal of Evolutionary Biology*, 15, 49-56.

Morimoto, J., Pizzari, T. & Wigby, S. (2016). Developmental environment effects on sexual selection in male and female *Drosophila melanogaster*. *PLoS One*, 11, 27.

Munroe, K.E. & Koprowski, J.L. (2011). Sociality, Bateman's gradients, and the polygynandrous genetic mating system of round-tailed ground squirrels (*Xerospermophilus tereticaudus*). *Behav. Ecol. Sociobiol.*, 65, 1811-1824.

Nakadera, Y., Marien, J., Van Straalen, N.M. & Koene, J.M. (2017). Multiple mating in natural populations of a simultaneous hermaphrodite, *Lymnaea stagnalis*. *Journal of Molluscan Studies*, 83, 56-62.

Paczolt, K.A., Passow, C.N., Delclos, P.J., Kindsvater, H.K., Jones, A.M.G. & Rosenthal, G.G. (2015). Multiple mating and reproductive skew in parental and introgressed females of the live-bearing fish *Xiphophorus birchmanni*. *J. Hered.*, 106, 57-66.

Pelissie, B., Jarne, P. & David, P. (2012). Sexual selection without sexual dimorphism: Bateman gradients in a simultaneous hermaphrodite. *Evolution; international journal of organic evolution*, 66, 66-81.

Poesel, A., Gibbs, H.L. & Nelson, D.A. (2011). Extrapair fertilizations and the potential for sexual selection in a socially monogamous songbird. *Auk*, 128, 770-776.

Pongratz, N. & Michiels, N.K. (2003). High multiple paternity and low last-male sperm precedence in a hermaphroditic planarian flatworm: consequences for reciprocity patterns. *Mol. Ecol.*, 12, 1425-1433.

Prosser, M.R., Weatherhead, P.J., Gibbs, H.L. & Brown, G.P. (2002). Genetic analysis of the mating system and opportunity for sexual selection in northern water snakes (*Nerodia sipedon*). *Behav. Ecol.*, 13, 800-807.

Ratkiewicz, M. & Borkowska, A. (2000). Multiple paternity in the bank vole (*Clethrionomys glareolus*): field and experimental data. *Zeitschrift Fur Saugetierkunde-International Journal of Mammalian Biology*, 65, 6-14.

Rios-Cardenas, O. (2005). Patterns of parental investment and sexual selection in teleost fishes: Do they support Bateman's principles? *Integrative and Comparative Biology*, 45, 885-894.

Rost, R. & Honegger, H.W. (1987). The timing of premating and mating behavior in a field population of the cricket *Gryllus campestris* L. *Behav. Ecol. Sociobiol.*, 21, 279-289.

Saunders, K.M. & Shuster, S.M. (2019). Bateman gradients and alternative mating strategies in a marine isopod. *IntechOpen*, (DOI: 10.5772/intechopen.88956).

Scheepers, M.J. & Gouws, G. (2019). Mating system, reproductive success, and sexual selection in Bluntnose Klipfishes (*Clinus cottoides*). *J. Hered.*, 110, 351-360.

Schlicht, E. & Kempenaers, B. (2013). Effects of social and extra-pair mating on sexual selection in blue tits (*Cyanistes caeruleus*) *Evolution; international journal of organic evolution*, 67, 1420-1434.

Schulte-Hostedde, A.I., Millar, J.S. & Gibbs, H.L. (2004). Sexual selection and mating patterns in a mammal with female-biased sexual size dimorphism. *Behav. Ecol.*, 15, 351-356.

Strausberger, B.M. & Ashley, M.V. (2003). Breeding biology of brood parasitic brown-headed cowbirds (*Molothrus ater*) characterized by parent-offspring and sibling-group reconstruction. *Auk*, 120, 433-445.
Tatarenkov, A., Healey, C.I.M., Grether, G.F. & Avise, J.C. (2008). Pronounced reproductive skew in a natural population of green swordtails, *Xiphophorus helleri*. *Mol. Ecol.*, 17, 4522-4534.
Turnell, B.R. & Shaw, K.L. (2015). High opportunity for postcopulatory sexual selection under field conditions. *Evolution; international journal of organic evolution*, 69, 2094-2104.
Ursprung, E., Ringler, M., Jehle, R. & Hodl, W. (2011). Strong male/male competition allows for nonchoosy females: high levels of polygynandry in a territorial frog with paternal care. *Mol. Ecol.*, 20, 1759-1771.
Walker, L.K., Ewen, J.G., Brekke, P. & Kilner, R.M. (2014). Sexually selected dichromatism in the hihi *Notiomystis cincta*: multiple colours for multiple receivers. *Journal of Evolutionary Biology*, 27, 1522-1535.
Wang, X., Liu, S., Yang, Y.Q., Wu, L.N., Huang, W.H., Wu, R.X. *et al.* (2020). Genetic evidence for the mating system and reproductive success of black sea bream (*Acanthopagrus schlegelii*). *Ecol. Evol.*, 10, 4483-4494.
Wells, C.P., Tomalty, K.M., Floyd, C.H., McElreath, M.B., May, B.P. & Van Vuren, D.H. (2017). Determinants of multiple paternity in a fluctuating population of ground squirrels. *Behav. Ecol. Sociobiol.*, 71, 13.
Whittingham, L.A. & Dunn, P.O. (2005). Effects of extra-pair and within-pair reproductive success on the opportunity for selection in birds. *Behav. Ecol.*, 16, 138-144.
Whittingham, L.A. & Lifjeld, J.T. (1995). High paternal investment in unrelated young: extra-pair paternity and male parental care in house martins. *Behav. Ecol. Sociobiol.*, 37, 103-108.

Supplementary Data

### List of primary studies

The list below encompasses all 77 published primary studies used in the meta-analysis on sexual selection in females. It does not include an unpublished study by Fromonteil et al. (in prep.).


Andrade, M.C.B. & Kasumovic, M.M. (2005). Terminal investment strategies and male mate choice: Extreme tests of Bateman. *Integrative and Comparative Biology*, 45, 838-847.

Anthes, N., David, P., Auld, J.R., Hoffer, J.N., Jarne, P., Koene, J.M. *et al.* (2010). Bateman gradients in hermaphrodites: an extended approach to quantify sexual selection. *Am. Nat.*, 176, 249-263.

Aronsen, T., Berglund, A., Mobley, K.B., Ratikainen, I.I. & Rosenqvist, G. (2013). Sex ratio and density affect sexual selection in a sex-role reversed fish. *Evolution*, 67, 3243-3257.

Balenger, S., Johnson, L. & Masters, B. (2009). Sexual selection in a socially monogamous bird: male color predicts paternity success in the mountain bluebird, *Sialia currucoides*. *Behav. Ecol. Sociobiol.*, 63, 403-411.

Barreto, F.S. & Avise, J.C. (2010). Quantitative measures of sexual selection reveal no evidence for sex-role reversal in a sea spider with prolonged paternal care. *Proceedings of the Royal Society B-Biological Sciences*, 277, 2951-2956.

Becher, S.A. & Magurran, A.E. (2004). Multiple mating and reproductive skew in Trinidadian guppies. *Proceedings of the Royal Society B-Biological Sciences*, 271, 1009-1014.

Bergeron, P., Montiglio, P.O., Reale, D., Humphries, M.M. & Garant, D. (2012). Bateman gradients in a promiscuous mating system. *Behav. Ecol. Sociobiol.*, 66, 1125-1130.

Bjork, A. & Pitnick, S. (2006). Intensity of sexual selection along the anisogamy-isogamy continuum. *Nature*, 441, 742-745.

Bolopo, D., Canestrari, D., Martinez, J.G., Roldan, M., Macias-Sanchez, E., Vila, M. *et al.* (2017). Flexible mating patterns in an obligate brood parasite. *Ibis*, 159, 103-112.

Borgerhoff Mulder, M. (2009). Serial monogamy as polygyny or polyandry? *Human Nature*, 20, 130-150.

Borgerhoff Mulder, M. & Ross, C.T. (2019). Unpacking mating success and testing Bateman's principles in a human population. *Proceedings of the Royal Society B-Biological Sciences*, 286, 10.

Broquet, T., Jaquiery, J. & Perrin, N. (2009). Opportunity for sexual selection and effective population size in the lek-breeding European treefrog (*Hyla arborea*) *Evolution*, 63, 674-683.

Burkli, A. & Jokela, J. (2017). Increase in multiple paternity across the reproductive lifespan in a sperm-storing, hermaphroditic freshwater snail. *Mol. Ecol.*, 26, 5264-5278.

Byers, B.E., Mays, H.L., Stewart, I.R.K. & Westneat, D.F. (2004). Extrapair paternity increases variability in male reproductive success in the chestnut-slded warbler (*Dendroica pensylvanica*), a socially monogamous songbird. *Auk*, 121, 788-795.

Cattelan, S., Evans, J.P., Garcia-Gonzalez, F., Morbiato, E. & Pilastro, A. (2020). Dietary stress increases the total opportunity for sexual selection and modifies selection on condition-dependent traits. *Ecol. Lett.*, 23, 447-456.

Collet, J., Richardson, D.S., Worley, K. & Pizzari, T. (2012). Sexual selection and the differential effect of polyandry. *Proc. Natl. Acad. Sci. U. S. A.*, 109, 8641-8645.

Courtiol, A., Pettay, J.E., Jokela, M., Rotkirch, A. & Lummaa, V. (2012). Natural and sexual selection in a monogamous historical human population. *Proc. Natl. Acad. Sci. U. S. A.*, 109, 8044-8049.

Croshaw, D.A. (2010). Quantifying sexual selection: a comparison of competing indices with mating system data from a terrestrially breeding salamander. *Biol. J. Linnean Soc.*, 99, 73-83.

Devost, E. & Turgeon, J. (2016). The combined effects of pre- and post-copulatory processes are masking sexual conflict over mating rate in *Gerris buenoi*. *Journal of Evolutionary Biology*, 29, 167-177.

Fitze, P.S. & Le Galliard, J.F. (2011). Inconsistency between different measures of sexual selection. *Am. Nat.*, 178, 256-268.

Fritzsche, K. & Arnqvist, G. (2013). Homage to Bateman: sex roles predict sex differences in sexual selection. *Evolution*, 67, 1926-1936.

Fuxjager, L., Wanzenbock, S., Ringler, E., Wegner, K.M., Ahnelt, H. & Shama, L.N.S. (2019). Within-generation and transgenerational plasticity of mate choice in oceanic stickleback under climate change. *Philos. Trans. R. Soc. B-Biol. Sci.*, 374, 12.

Gagnon, M.-C., Duchesne, P. & Turgeon, J. (2012). Sexual conflict in *Gerris gillettei* (Insecta: Hemiptera): influence of effective mating rate and morphology on reproductive success. *Canadian Journal of Zoology*, 90, 1297-1306.

Garcia-Navas, V., Ferrer, E.S., Bueno-Enciso, J., Barrientos, R., Sanz, J.J. & Ortego, J. (2014). Extrapair paternity in Mediterranean blue tits: socioecological factors and the opportunity for sexual selection. *Behavioral Ecology*, 25, 228-238.

Gerlach, N.M., McGlothlin, J.W., Parker, P.G. & Ketterson, E.D. (2012). Reinterpreting Bateman gradients: multiple mating and selection in both sexes of a songbird species. *Behavioral Ecology*, 23, 1078-1088.

Glaudas, X., Rice, S.E., Clark, R.W. & Alexander, G.J. (2020). The intensity of sexual selection, body size and reproductive success in a mating system with male-male combat: is bigger better? *Oikos*, 129, 998-1011.

Gopurenko, D., Williams, R.N. & DeWoody, J.A. (2007). Reproductive and mating success in the small-mouthed salamander (*Ambystoma texanum*) estimated via microsatellite parentage analysis. *Evolutionary Biology*, 34, 130-139.

Gopurenko, D., Williams, R.N., McCormick, C.R. & DeWoody, J.A. (2006). Insights into the mating habits of the tiger salamander (*Ambystoma tigrinum tigrinum*) as revealed by genetic parentage analyses. *Mol. Ecol.*, 15, 1917-1928.

Grunst, A.S., Grunst, M.L., Korody, M.L., Forrette, L.M., Gonser, R.A. & Tuttle, E.M. (2019). Extrapair mating and the strength of sexual selection: insights from a polymorphic species. *Behavioral Ecology*, 30, 278-290.

Hoffer, J.N.A., Marien, J., Ellers, J. & Koene, J.M. (2017). Sexual selection gradients change over time in a simultaneous hermaphrodite. *eLife*, 6, 16.

Janicke, T., David, P. & Chapuis, E. (2015). Environment-dependent sexual selection: Bateman's parameters under varying levels of food availability. *Am. Nat.*, 185, 756-768.

Johannesson, K., Saltin, S.H., Charrier, G., Ring, A.K., Kvarnemo, C., Andre, C. *et al.* (2016). Non-random paternity of offspring in a highly promiscuous marine snail suggests postcopulatory sexual selection. *Behav. Ecol. Sociobiol.*, 70, 1357-1366.

Jokela, M., Rotkirch, A., Rickard, I.J., Pettay, J. & Lummaa, V. (2010). Serial monogamy increases reproductive success in men but not in women. *Behavioral Ecology*, 21, 906-912.

Jones, A.G., Arguello, J.R. & Arnold, S.J. (2002). Validation of Bateman's principles: a genetic study of sexual selection and mating patterns in the rough-skinned newt. *Proc. R. Soc. Lond. Ser. B-Biol. Sci.*, 269, 2533-2539.

Jones, A.G., Arguello, J.R. & Arnold, S.J. (2004). Molecular parentage analysis in experimental newt populations: The response of mating system measures to variation in the operational sex ratio. *Am. Nat.*, 164, 444-456.

Jones, A.G., Rosenqvist, G., Berglund, A., Arnold, S.J. & Avise, J.C. (2000). The Bateman gradient and the cause of sexual selection in a sex-role-reversed pipefish. *Proc. R. Soc. Lond. Ser. B-Biol. Sci.*, 267, 677-680.

Jones, P.H., Van Zant, J.L. & Dobson, F.S. (2012). Variation in reproductive success of male and female Columbian ground squirrels (*Urocitellus columbianus*). *Can. J. Zool.-Rev. Can. Zool.*, 90, 736-743.

Ketterson, E.D., Parker, P.G., Raouf, S.A., Nolan Jr, V., Ziegenfus, C. & Chandler, C.H. (1997). The relative impact of extra-pair fertilizations on variation in male and female reproductive success in dark-eyed juncos (*Junco hyemaus*). In: *Avian Reproductive Tactics: Female and Male Perspectives* (eds. Parker, PG & Burley, NT), pp. 81-101.

Krakauer, A.H. (2008). Sexual selection and the genetic mating system of Wild Turkeys. *Condor*, 110, 1-12.

Kretzschmar, P., Auld, H., Boag, P., Ganslosser, U., Scott, C., de Groot, P.J.V. *et al.* (2020). Mate choice, reproductive success and inbreeding in white rhinoceros: New insights for conservation management. *Evol. Appl.*, 13, 699-714.

LaBrecque, J.R., Alva-Campbell, Y.R., Archambeault, S. & Crow, K.D. (2014). Multiple paternity is a shared reproductive strategy in the live-bearing surfperches (Embiotocidae) that may be associated with female fitness. *Ecol. Evol.*, 4, 2316-2329.

Levine, B.A., Schuett, G.W., Clark, R.W., Repp, R.A., Herrmann, H.W. & Booth, W. (2020). No evidence of male-biased sexual selection in a snake with conventional Darwinian sex roles. *R. Soc. Open Sci.*, 7, 10.

Levine, B.A., Smith, C.F., Schuett, G.W., Douglas, M.R., Davis, M.A. & Douglas, M.E. (2015). Bateman-Trivers in the 21st Century: sexual selection in a North American pitviper. *Biol. J. Linnean Soc.*, 114, 436-445.

Levitan, D.R. (2008). Gamete traits influence the variance in reproductive success, the intensity of sexual selection, and the outcome of sexual conflict among congeneric sea urchins. *Evolution*, 62, 1305-1316.

Louder, M.I.M., Hauber, M.E., Louder, A.N.A., Hoover, J.P. & Schelsky, W.M. (2019). Greater opportunities for sexual selection in male than in female obligate brood parasitic birds. *Journal of Evolutionary Biology*, 32, 1310-1315.

Mangold, A., Trenkwalder, K., Ringler, M., Hoedl, W. & Ringler, E. (2015). Low reproductive skew despite high male-biased operational sex ratio in a glass frog with paternal care. *Bmc Evolutionary Biology*, 15.

Marie-Orleach, L., Janicke, T., Vizoso, D.B., David, P. & Scharer, L. (2016). Quantifying episodes of sexual selection: Insights from a transparent worm with fluorescent sperm. *Evolution*, 70, 314-328.

McCullough, E.L., Buzatto, B.A. & Simmons, L.W. (2018). Population density mediates the interaction between pre- and postmating sexual selection. *Evolution*, 72, 893-905.

Mills, S.C., Grapputo, A., Koskela, E. & Mappes, T. (2007). Quantitative measure of sexual selection with respect to the operational sex ratio: a comparison of selection indices. *Proceedings of the Royal Society B-Biological Sciences*, 274, 143-150.

Mobley, K.B. & Jones, A.G. (2013). Overcoming statistical bias to estimate genetic mating systems in open populations: a comparison of Bateman's principles between the sexes in a sex-role-reversed pipefish. *Evolution*, 67, 646-660.

Moorad, J.A., Promislow, D.E.L., Smith, K.R. & Wade, M.J. (2011). Mating system change reduces the strength of sexual selection in an American frontier population of the 19th century. *Evolution and Human Behavior*, 32, 147-155.

Morimoto, J., Pizzari, T. & Wigby, S. (2016). Developmental environment effects on sexual selection in male and female *Drosophila melanogaster*. *PLoS One*, 11, 27.

Munroe, K.E. & Koprowski, J.L. (2011). Sociality, Bateman's gradients, and the polygynandrous genetic mating system of round-tailed ground squirrels (*Xerospermophilus tereticaudus*). *Behav. Ecol. Sociobiol.*, 65, 1811-1824.

Paczolt, K.A., Passow, C.N., Delclos, P.J., Kindsvater, H.K., Jones, A.M.G. & Rosenthal, G.G. (2015). Multiple mating and reproductive skew in parental and introgressed females of the live-bearing fish *Xiphophorus birchmanni*. *J. Hered.*, 106, 57-66.

Pelissie, B., Jarne, P. & David, P. (2012). Sexual selection without sexual dimorphism: Bateman gradients in a simultaneous hermaphrodite. *Evolution*, 66, 66-81.

Poesel, A., Gibbs, H.L. & Nelson, D.A. (2011). Extrapair fertilizations and the potential for sexual selection in a socially monogamous songbird. *Auk*, 128, 770-776.

Pongratz, N. & Michiels, N.K. (2003). High multiple paternity and low last-male sperm precedence in a hermaphroditic planarian flatworm: consequences for reciprocity patterns. *Mol. Ecol.*, 12, 1425-1433.

Prosser, M.R., Weatherhead, P.J., Gibbs, H.L. & Brown, G.P. (2002). Genetic analysis of the mating system and opportunity for sexual selection in northern water snakes (*Nerodia sipedon*). *Behavioral Ecology*, 13, 800-807.

Rios-Cardenas, O. (2005). Patterns of parental investment and sexual selection in teleost fishes: Do they support Bateman's principles? *Integrative and Comparative Biology*, 45, 885-894.

Rodriguez-Munoz, R., Bretman, A., Slate, J., Walling, C.A. & Tregenza, T. (2010). Natural and sexual selection in a wild insect population. *Science*, 328, 1269-1272.

Rose, E., Paczolt, K.A. & Jones, A.G. (2013). The contributions of premating and postmating selection episodes to total selection in sex-role-reversed Gulf Pipefish. *Am. Nat.*, 182, 410-420.

Saunders, K.M. & Shuster, S.M. (2019). Bateman gradients and alternative mating strategies in a marine isopod. *IntechOpen*, (DOI: 10.5772/intechopen.88956).

Scheepers, M.J. & Gouws, G. (2019). Mating system, reproductive success, and sexual selection in Bluntnose Klipfishes (*Clinus cottoides*). *J. Hered.*, 110, 351-360.

Schlicht, E. & Kempenaers, B. (2013). Effects of social and extra-pair mating on sexual selection in blue tits (*Cyanistes caeruleus*) *Evolution*, 67, 1420-1434.

Schulte-Hostedde, A.I., Millar, J.S. & Gibbs, H.L. (2004). Sexual selection and mating patterns in a mammal with female-biased sexual size dimorphism. *Behavioral Ecology*, 15, 351-356.

Serbezov, D., Bernatchez, L., Olsen, E.M. & Vollestad, L.A. (2010). Mating patterns and determinants of individual reproductive success in brown trout (*Salmo trutta*) revealed by parentage analysis of an entire stream living population. *Mol. Ecol.*, 19, 3193-3205.

Skjaervo, G.R. & Roskaft, E. (2015). Wealth and the opportunity for sexual selection in men and women. *Behavioral Ecology*, 26, 444-451.

Tatarenkov, A., Healey, C.I.M., Grether, G.F. & Avise, J.C. (2008). Pronounced reproductive skew in a natural population of green swordtails, *Xiphophorus helleri*. *Mol. Ecol.*, 17, 4522-4534.

Turnell, B.R. & Shaw, K.L. (2015). High opportunity for postcopulatory sexual selection under field conditions. *Evolution*, 69, 2094-2104.

Ursprung, E., Ringler, M., Jehle, R. & Hodl, W. (2011). Strong male/male competition allows for nonchoosy females: high levels of polygynandry in a territorial frog with paternal care. *Mol. Ecol.*, 20, 1759-1771.

Walker, L.K., Ewen, J.G., Brekke, P. & Kilner, R.M. (2014). Sexually selected dichromatism in the hihi *Notiomystis cincta*: multiple colours for multiple receivers. *Journal of Evolutionary Biology*, 27, 1522-1535.

Wang, X., Liu, S., Yang, Y.Q., Wu, L.N., Huang, W.H., Wu, R.X. *et al.* (2020). Genetic evidence for the mating system and reproductive success of black sea bream (*Acanthopagrus schlegelii*). *Ecol. Evol.*, 10, 4483-4494.

Wells, C.P., Tomalty, K.M., Floyd, C.H., McElreath, M.B., May, B.P. & Van Vuren, D.H. (2017). Determinants of multiple paternity in a fluctuating population of ground squirrels. *Behav. Ecol. Sociobiol.*, 71, 13.

Whittingham, L.A. & Dunn, P.O. (2005). Effects of extra-pair and within-pair reproductive success on the opportunity for selection in birds. *Behavioral Ecology*, 16, 138-144.

Whittingham, L.A. & Lifjeld, J.T. (1995). High paternal investment in unrelated young: extra-pair paternity and male parental care in house martins. *Behav. Ecol. Sociobiol.*, 37, 103-108.

Williams, R.N. & DeWoody, J.A. (2009). Reproductive success and sexual selection in wild eastern tiger salamanders (*Ambystoma t. tigrinum*). *Evolutionary Biology*, 36, 201-213.

Woolfenden, B.E., Gibbs, H.L. & Sealy, S.G. (2002). High opportunity for sexual selection in both sexes of an obligate brood parasitic bird, the brown-headed cowbird (*Molothrus ater*). *Behav. Ecol. Sociobiol.*, 52, 417-425.
